# Mapping cherry blossoms from geotagged street-level photos

**DOI:** 10.1101/2022.01.18.476550

**Authors:** Shuya Funada, Narumasa Tsutsumida

## Abstract

Floral phenology is useful information as an indicator on climate change and ecosystem services, however its observation is not straightforward over space and time. Satellite remote sensing and official and volunteer-based *in-situ* observations have been conducting, but the long-term and accurate data collection is challenging due to the insufficient quality and quantity of observations and the lack of financial and human resources to sustain. Here, we demonstrate a flower detection model from street-level photos, which can be the core function of a semi-automatic observation system to tackle those issues above. We detected cherry blossoms by this model from geotagged images with the observation date, obtained from Mapillary, which is one of social sensing data sources, and mapped dates of flowering in a study site, Aizuwakamatsu, Japan in April 2018. This approach enables us to collect floral phenology information semi-automatically as a data-driven approach. It is expected to collect a large number of observations with a certain level of quality by avoiding human-induced biases for the observations.

## 1. INTRODUCTION

Monitoring floral phenology is challenging. Many satellite-based remote sensing techniques have been applied to floral phenological studies and some successful approaches has been reported [1], [2]; however, it is not yet straightforward to identify flowering events due to less observation frequency and coarse spatial resolution. The long-term *in-situ* observation has been making efforts to record phenology. For example, Japan Meteorological Agency (JMA) has been conducting seasonal observations of plants and animals every year since 1953 to monitor seasonal changes in their condition. However, the target has significantly been declined since 2021 because of difficulties observing some species, with inadequate funding [3]. The 40% of target sites have been terminated, and 94% of phenological events (including plants and animals) have been no more observed. Instead, citizen science programs attempt to substitute the phenological observations. Although we expect the quantity and quality of observations to be the same as JMA did, it is still challenging to implement citizen science approaches robustly because of human-related issues. Not all observers are not trained enough to monitor phenological events, and thus records collected from observers would include some inevitable errors and biases [4], [5]. As such human-induced issues may be critical obstacles for phenological monitoring, a semi-automatic observation system would be helpful to collect and record phenological events over space and time by a data-driven approach. To achieve this, we demonstrate a flower detection model from street-level photos, which can be the core function of such a system. We focus on cherry blossoms in spring as a first step because the flowering cherry blossoms can be an important indicator of environmental change and cultural ecosystem services [6]. Our model enables us to detect cherry blossoms from a street-level photo, and as such, it is applicable to social sensing photo data with the observation date and geolocation to map cherry blossoms. We obtained a series of street-level photos via Mapillary API in Aizuwakamatsu, Japan, in April 2019 and successfully recorded flowering events over space and time.

Although cherry blossoms will be monitored continuously by JMA, our approach can be an additional way to record flowering events spatiotemporally.

## 2. MATERIALS AND METHODS

We build a deep-learning model by ‘YOLOv4’ [7] to detect cherry blossoms from a street-level photo. In the total of 200 photos are annotated manually to train cherry flowers (Figure 1). We only target Japanese cherry trees (*Prunus* × *yedoensis*, Someiyoshino) because this species is the most popular and widely planted across Japan. Estimated photos with a detection probability of over 80% are only selected as cherry blossoms.

**Figure 1.**
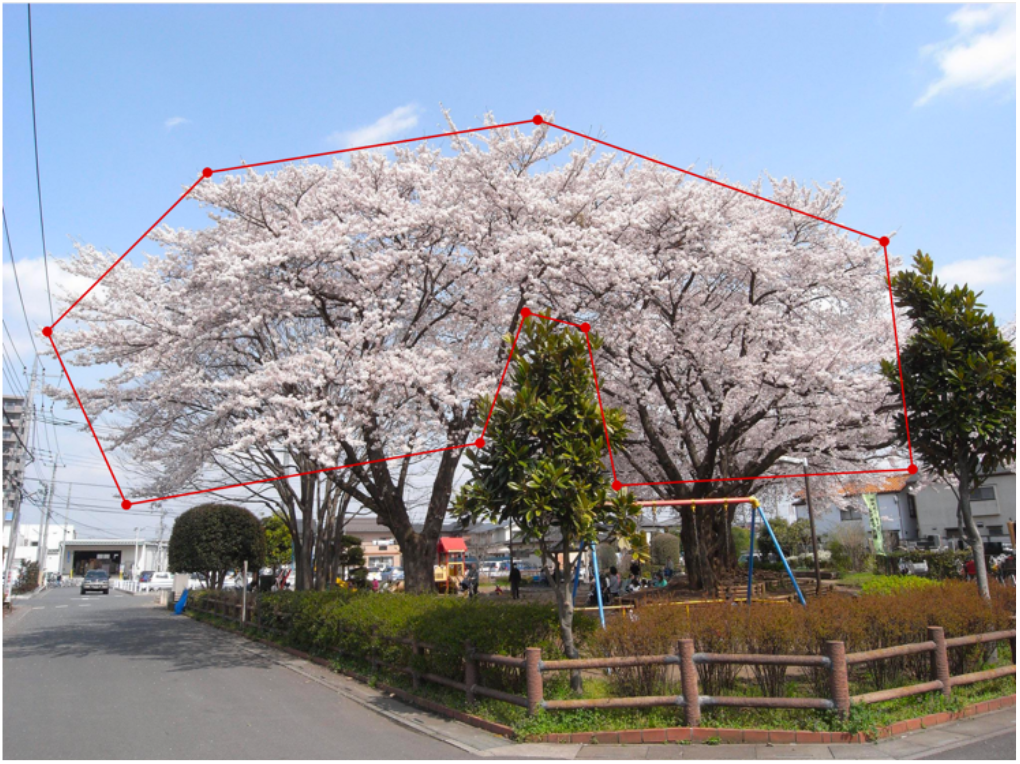
An example of images with the annotation of cherry blossoms.

We set the study area in Aizuwakamatsu, Japan, covering famous Cherry blossom sightseeing spots, Tsurugajo castle and Ishibe-Sakura. We collect 28,999 street-level photos via Mapillary (https://www.mapillary.com/) within the study area from 1^st^ April to 30^th^ April 2018. The target period is overlapped with the first start flowering date in 2018 (4^th^ April) at Tsurugajo castle, according to the report found at https://www.tsurugajo.com/turugajo/sakura.html. 2,570 of estimated photos were validated by visual inspection to evaluate whether cherry blossoms were successfully detected or not.

## 3. RESULTS

A successful example of the detection of the cherry blossoms in a street-level photo is shown in Figure 2. Our model estimates the extent of flowers on a Japanese cherry tree and does not attempt to detect every flower in a photo. By using this approach, we make a cherry blossom map from Mapillary photos (Figure 3). This indicates that cherry blossom events vary over space and time. It is yet difficult to continuously monitoring due to insufficient photos available, but this map depicts when and where cherry blossoms were found.

**Figure 2.**
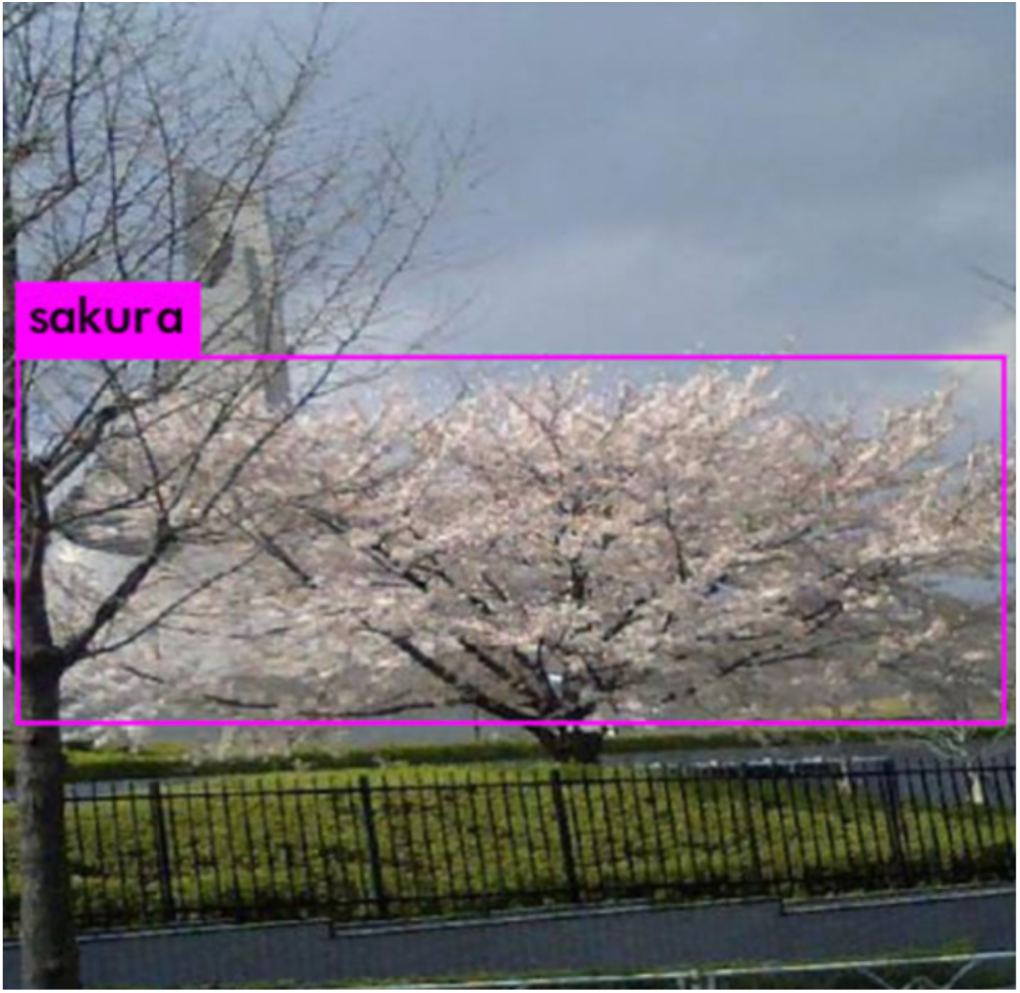
An example of the cherry blossom detection from a street-level photo by our model.

**Table 1.** A confusion matrix of the detection result by the proposed model from 2,570 samples.

|  |  | Model results |  |
| --- | --- | --- | --- |
|  |  | Detected | undetected |
| Reference | Positive | 371 | 350 |
|  | Negative | 90 | 1759 |

**Figure3.**
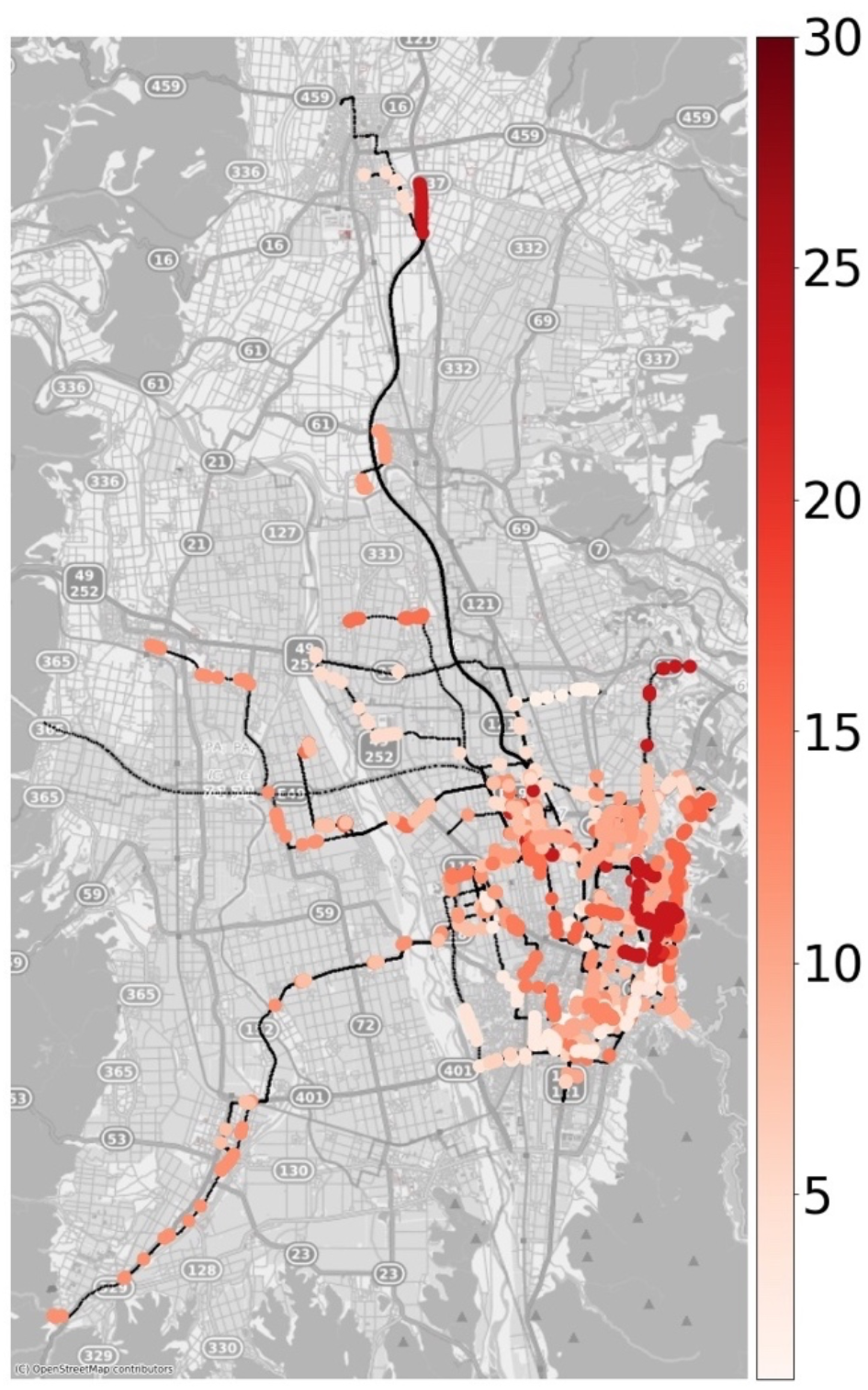
A result cherry blossoms detections. The legend denotes days in April. Black dots represent no cherry blossoms found in photos.

The confusion matrix reports the evaluation of the detection from 2,570 test photos. The overall accuracy was 0.83. Precision and recall were 0.80 and 0.51, respectively. Low recall suggests that the detection model may tend to overlook the actual flowering events.

## 4. DISCUSSION

The object-detection model we built detects the cherry blossom from street-level photos with 83% of accuracy. This achievement contributes to the development of a semi-automatic phenological observation system using social sensing photo data. The detection ability depends on the model performance. Our model shows relatively low recall, suggesting the result may tend to overlook cherry blossoms found in a photo to some extent. This issue should be overcome to develop the model with additional trained photos by annotation efforts. However, as the bias of misclassification relies on the model performance, we expect to minimize human errors and biases for observations. The critical issue of this approach is the data availability. A large number of photos obtained via Mapillary are available in the study area and the study period, but the frequency and spatial extent of observations are not sufficient to observe all floral phenological events. Thus, it should be noted that the map shown in Figure 3 represents not the timing of flowering events, but the timing of the flower detection.

## 5. CONCLUSIONS

We propose a novel approach to detect cherry blossoms from street-level photos. The object-detection model we built achieved 83% overall accuracy: relatively high precision of 80% but low recall of 52%. This model was applied to social sensing photos sourced from Mapillary. The cherry blossom map tells when and where flowering was found. This approach is a novel semi-automatic observation as a data-driven approach. Further developments on the model will be needed to reduce the misclassification rate. In the future, we also attempt to develop models for other species to extract phenological information from a set of available data sources.

## ACKNOWLEDGEMENTS

We appreciate Mapillary users to share valuable street-level photos. This work is partly supported by National Geographic Asia Labs Fellow.

## Notes

### Competing Interest Statement

The authors have declared no competing interest.

